# A lectin receptor-like kinase controls self-pollen recognition in *Phlox* (Polemoniaceae)

**DOI:** 10.1101/2025.06.17.660178

**Authors:** Grace A. Burgin, Nia Lewis, Robin Hopkins

## Abstract

Self-incompatibility (SI) describes a widespread collection of genetic mechanisms in flowering plants used to specifically recognize and reject self-pollen. These mechanisms are fundamental to plant sexual reproduction and offer valuable insight into the molecular basis of cell-cell communication and self-recognition more broadly. Here, we leverage an independent evolution of SI in the lineage containing *Phlox* (Polemoniaceae) to characterize a novel gene causing self-pollen recognition which we name *Phlox drummondii* Pistil Identity Receptor Kinase (*PdPIRK*). Recognition of self-pollen associates with a single genomic region containing the *Phlox S*-locus. We generate predictions regarding how *S*-loci must function and evolve to identify a single candidate gene within this *S*-associated region. This gene, *PdPIRK*, is highly and specifically expressed in the pistil and has exceptionally high polymorphism maintained by negative frequency dependent selection, two hallmarks of self-pollen recognition genes. Functional validation with gene silencing confirms that *PdPIRK* is necessary for self-incompatibility, and we further demonstrate allele specific activity, confirming its role in self-pollen recognition per se. *PdPIRK* encodes a G-type lectin receptor-like kinase, which is a member of the same gene family as *SRK*, the gene controlling self-pollen recognition in the distantly related Brassicaceae. Our findings suggest the presence of genetic constraints or paths of least resistance governing how *S*-loci evolve and add to our understanding of the diverse molecular mechanisms through which organisms achieve self-recognition.

## Introduction

Distinguishing self from non-self is a fundamental feature of many biological processes. Self-avoidance drives neuronal geometries in insects and vertebrates, a breakdown in self-tolerance leads to aberrant immune phenotypes in mammals, and the spatial boundaries of an expanding bacterial colony can be determined through encounters with non-self-strains^1–3^. Identifying the genetic basis of these self-discriminating processes has contributed essential insight into how molecular communication functions and evolves. However, many self-recognition mechanisms remain to be discovered, particularly across diverse and non-model taxa.

In flowering plants, reliable self- and non-self-discrimination is essential during sexual reproduction. Most angiosperms produce sperm-bearing pollen and ovule-housing pistils within the same individual which generates the potential for self-fertilization^4, 5^. The rate of self-fertilization can affect individual fitness, recombination dynamics, and how evolution proceeds across the entire genome^6^. As a result, plants have evolved diverse mechanisms to modulate the rate of self-fertilization within a lineage. The most widespread of these mechanisms is self-incompatibility (SI) which describes genetic mechanisms used to specifically recognize and reject self-pollen when it arrives at the pistil. SI independently evolved numerous times as flowering plants diversified, and the general term of SI refers collectively to these varied origins^7, 8^. In all cases, functional SI relies on recognition of self-pollen via molecular interactions between pistil and pollen expressed gene products encoded within a single, non-recombining genomic locus termed the *S*-locus^9, 10^. These systems most commonly involve many compatibility types determined by an individual’s *S*-locus genotype without associated variation in floral morphology (i.e., homomorphic SI).

To date, the genetic identities of just five independently evolved homomorphic *S*-loci are known and each encodes unique and nonhomologous genes^11–15^. This observation suggests that self-pollen recognition can and does evolve through diverse genetic routes. Whether this apparent nonhomology reflects a true lability in how self-pollen recognition functions and evolves is unclear because the majority of independently evolved *S*-loci remain uncharacterized^7^.

*Phlox drummondii* (Polemoniaceae) presents a unique opportunity to study the evolution of self-pollen recognition. *Phlox* is a classic system for exploring evolutionary transitions in the SI response, yet the genes causing self-pollen recognition and rejection remain unknown^16, 17^. Its phylogenetic position indicates that an independent evolution of SI has occurred within the lineage containing *Phlox* (Figure 1A)^18^. We leverage this independent evolution to characterize the genetic identity of the pistil-expressed gene of a novel *S*-locus^19^. Our study adds fundamental insight into the diverse molecular mechanisms through which organisms achieve self-discrimination.

**Figure 1:**
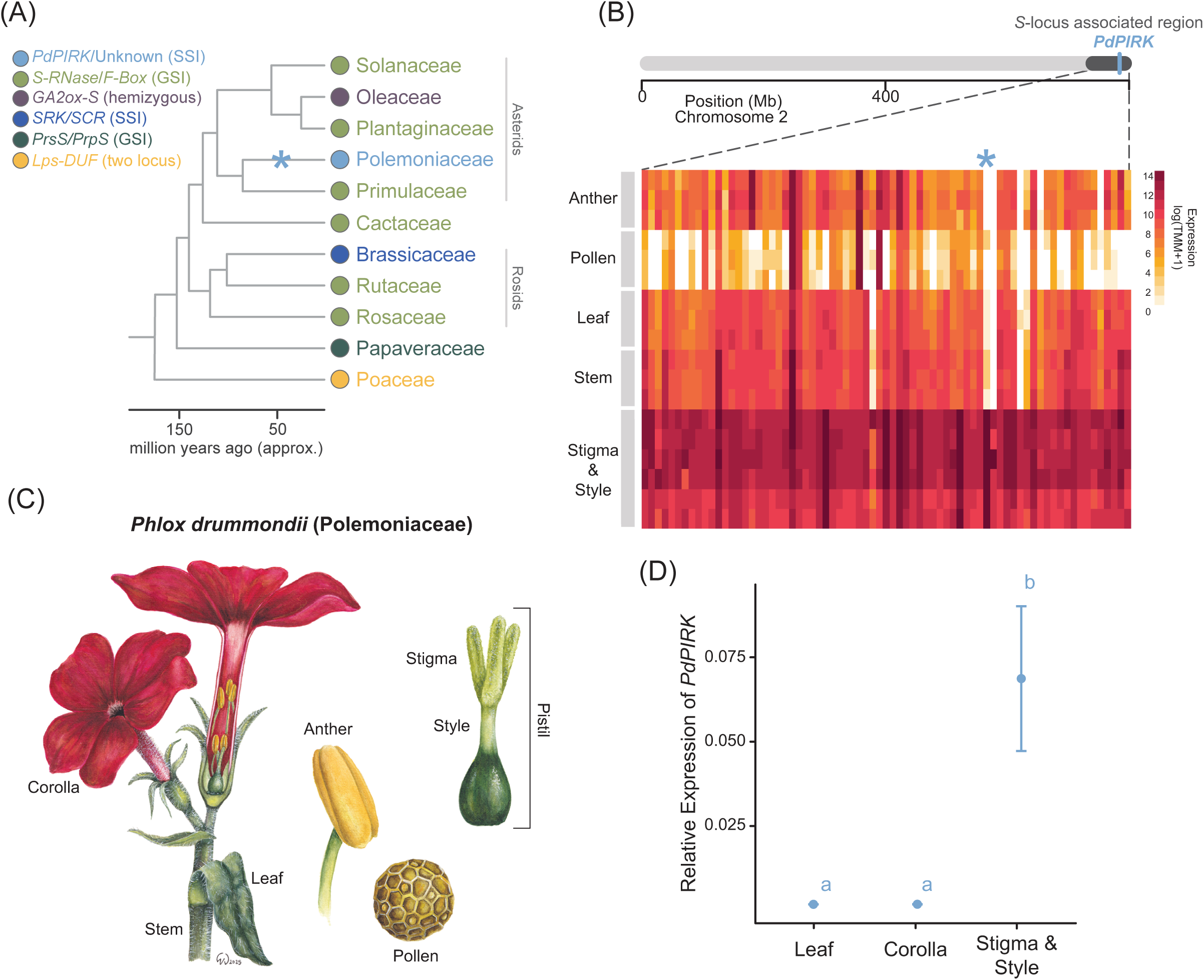
Identifying a candidate gene for self-pollen recognition in *P. drummondii.* **(A)** A schematic of the distribution of independently evolved homomorphic SI mechanisms for which the genetic identity of the *S*-locus is known across angiosperms. Colors indicate unique SI mechanisms found in each family with the pistil/pollen expressed *S*-locus genetic components listed in the legend. The categorical SI type is indicated in parentheses: sporophytic given as SSI, gametophytic as GSI, or two locus system^22^. An independent evolution of SI has occurred in the lineage containing *Phlox* (Polemoniaceae) and is indicated by a blue asterisk. **(B)** Schematic of *P. drummondii* Chromosome 2 with dark gray shading indicating the ∼67Mb region our previous genetic mapping experiments identified as associated with the *S*-locus. The approximate genomic position of *PdPIRK* is marked by a blue line. The top ∼10% of genes (n=73) with highest expression in stigma/style tissue in this genomic region are shown below in a heatmap of expression with tissue type and biological replicates organized into rows (Supplemental Table S1&S2). Each column is a unique gene and genes are ordered based on their physical position in the genome. A blue asterisk denotes *PdPIRK* which is highly and specifically expressed in stigma/style tissue. **(C)** Illustration of *P. drummondii* floral morphology by Dr. Wendy Van Drunen (www.evoecoart.com) with tissues included in transcriptome sequencing and/or qPCR experiments labelled. **(D)** The relative expression of *PdPIRK* in leaf, corolla, and stigma/style tissue from *S4S4* plants as quantified by qPCR corroborates a pattern of stigma/style specific expression. Primers used to amplify *PdPIRK* were based on the allele *S4* sequence identified from long-read transcriptomes (Supplemental Table S3). Means ± standard errors are plotted with lower case letters indicating significance differences between tissue types.

## Results

### A single pistil-expressed gene matches predictions for *S*-locus evolution

Self-pollen recognition in *P. drummondii* localizes to a single genomic locus spanning ∼67Mb and containing 727 total genes^18^. Here, we narrow this list of candidate genes using theoretical predictions for how self-pollen recognition genes must function and evolve. Specifically, we hypothesize that key features of the pistil-expressed *S*-locus gene include 1) high expression restricted to stigma/style tissue and 2) exceptionally high levels of polymorphism maintained by negative frequency-dependent selection.

Characterization of other self-pollen recognition mechanisms has revealed that while many genes are expressed in the pistil, the pistil-expressed *S*-locus gene is consistently among the most highly expressed in this tissue and rarely expressed elsewhere in the plant^20–23^. To identify genes following this expression pattern, we analyzed previously published transcriptomes of five *Phlox* tissues: stem, leaf, pollen, anthers, and stigma/style (Figure 1B&C). Tests for differential expression reveal that 224 genes genome-wide show significantly higher expression in the stigma/style in pairwise contrasts with all other tissue types. Of the 727 genes annotated to the *S*-locus associated genomic region, eight are restricted in expression to stigma/style tissue (Supplemental Table S1). We validated the pattern of tissue specific expression for our top candidate gene in this region using quantitative real-time PCR (qPCR) (Figure 1D).

Negative frequency dependent selection maintains unusually high polymorphism at the *S*-locus^24^. In addition to not being able to self-fertilize, individuals with matching *S*-genotypes cannot fertilize each other. Rare alleles therefore confer a selective advantage over common alleles by increasing an individual’s pool of potential mating partners resulting in the maintenance of numerous alleles over evolutionary time and exceptionally high levels of polymorphism at genes within the *S*-locus. We quantify polymorphism across genes in the *S*-locus associated genomic region by calling phased variants using our stigma/style RNAseq dataset and reconstructing evolutionary relationships between the 12 alleles using maximum likelihood gene trees. Gene tree branch length is a metric for polymorphism with longer total branch lengths indicating higher overall polymorphism. Predicted coding regions of the top ∼10% of genes most highly expressed in stigma/style tissue within the *S*-locus associated genomic region have a mean ± SE total branch length of 0.0472 ± 0.0061 substitutions per site (n=73). There is a single outlier gene with an exceptionally long total branch length of 0.296 substitutions per site (Rosner’s Outlier Test R=4.75, k=1, p<0.001) (Figure 2A). Diversity at this gene is consistent with the extraordinarily high allelic variation predicted for functional genes within an *S*-locus.

**Figure 2:**
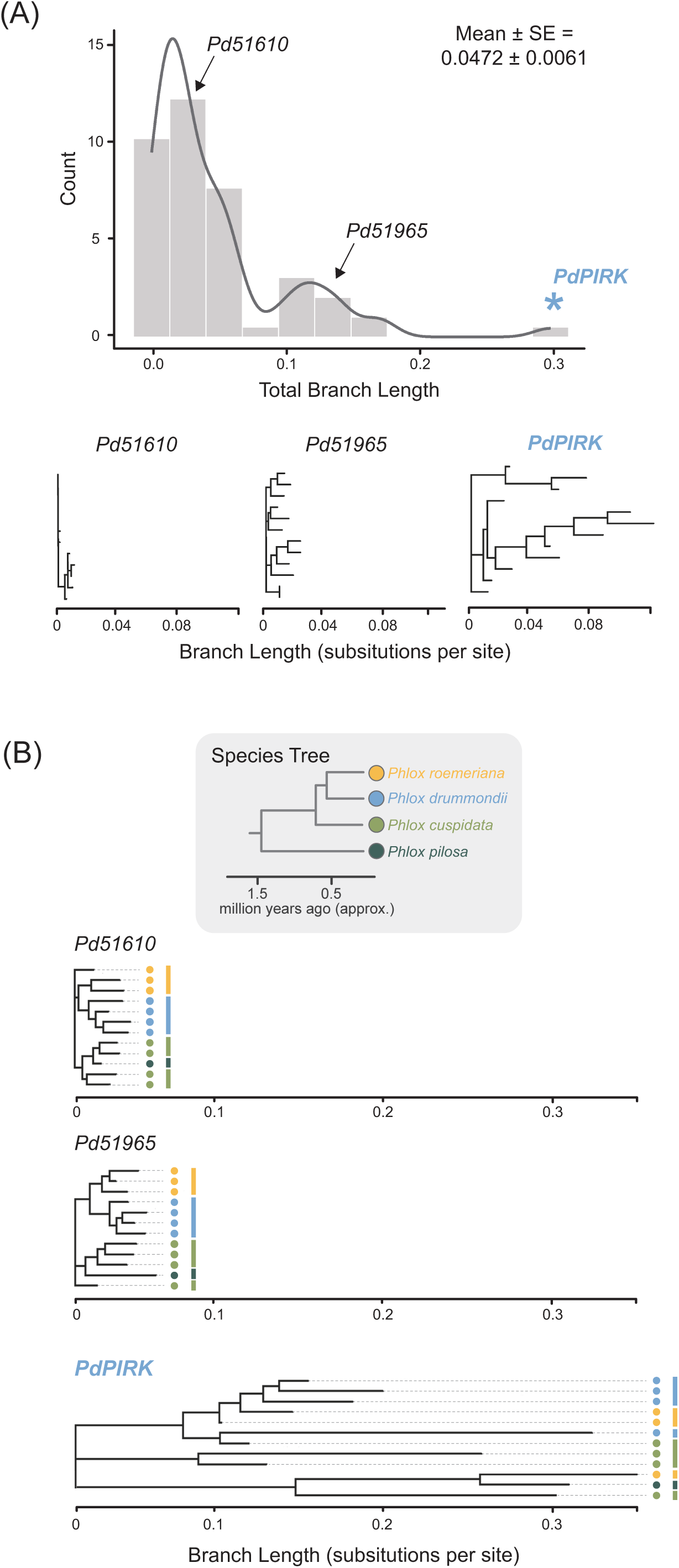
*PdPIRK* displays extreme polymorphism consistent with a signature of negative frequency dependent selection. **(A)** The distribution of total branch length (substitutions per site) for the top ∼10% most highly expressed genes in stigma/style tissue within the *S*-locus associated region. Gene trees were constructed using within-species allele sequences from *P. drummondii* individuals. The distribution contains a single outlier, *PdPIRK*, which is indicated by a blue asterisk. Gene trees of *PdPIRK* and two other genes are plotted along a standard axis of evolutionary change to exemplify branch length variation. **(B)** A schematic of the species relationships between four closely related *Phlox* species. Multi-species gene trees of *PdPIRK* and two other genes are plotted along a standard axis of evolutionary change with tips colored by species identity (*P. drummondii* in blue, *P. roemeriana* in yellow, *P. cuspidata* in light green, and *P. pilosa* in dark green). *PdPIRK* displays persistent allele sharing across species splits.

An additional signature of negative frequency dependent selection is trans-specific polymorphism (TSP), which describes persistent sharing of alleles across species splits due to common ancestry^25, 26^. To characterize TSP, we called variants in candidate genes within the *S*-locus associated genomic region from long-read sequencing data to construct multiple alleles sampled from each of four closely related *Phlox* species. We generated maximum likelihood gene trees and find that the gene with exceptionally long branch lengths within *P. drummondii* demonstrates persistent allele sharing between related *Phlox* species consistent with a pattern of TSP (Figure 2B).

Within the genomic region containing the *Phlox S*-locus, a single gene matches both predictions of 1) high and restricted stigma/style expression and 2) exceptional polymorphism as demonstrated by two independent analyses. We therefore hypothesize that this gene, which we name *P. drummondii* Pistil Identity Receptor Kinase (*PdPIRK)*, controls self-pollen recognition in *Phlox*.

### Genotypic variation at *PdPIRK* predicts cross-compatibility

We test the prediction that *PdPIRK* causes self-pollen recognition by confirming that genotypic variation predicts crossing success among individuals with shared alleles. Specifically, we expect that crosses between individuals with the same genotype will be recognized as self and produce no fruit while crosses between individuals with different genotypes will successfully produce fruit. To test this prediction, we crossed two unrelated individuals to generate ten full-sibling offspring each with one of four possible *S*-genotypes (Figure 3A). In our full diallel cross, we recover four groups of plants with near-identical crossing phenotypes corresponding to each of these four segregating *S*-genotypes (Figure 3B). Our previous crossing experiments reveal that *Phlox* SI is sporophytic, meaning that pollen self-identity is encoded by the diploid genotype of the anther-bearing plant in which the pollen developed rather than the haploid genotype of each pollen grain^18^. Allelic dominance is a common feature of sporophytic SI, and we observe a dominance hierarchy in which one allele is recessive to the other three co-dominant alleles captured in this cross (Figure 3A)^27^.

**Figure 3:**
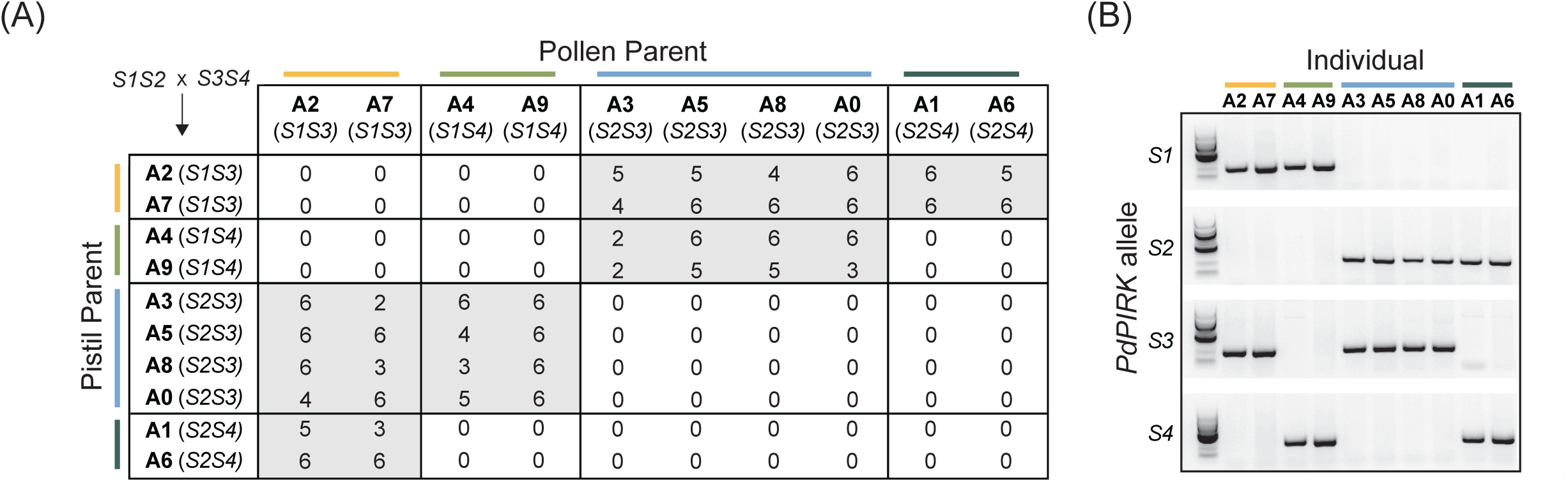
*PdPIRK* genotype predicts cross-compatibility phenotype. **(A)** A cross between an *S1S2* individual and *S3S4* individual generated ten full siblings with four possible *S*-locus genotypes. Results of a full diallel cross among siblings recovers four cross-compatibility groups, consistent with four *S*-locus genotypes segregating. Individuals are listed by their ID enumerated A0-A9. Inferred *S*-locus genotype based on cross-compatibility is given in parentheses. Each individual served as both a pistil parent (rows) and pollen parent (columns) for all possible cross combinations. The resulting number of fruits for each cross (out of six possible) is given and shaded based on compatibility, with compatible crosses shaded gray. As is common in systems with sporophytic SI, we recover dominance interactions between alleles. In both pollen and pistil tissues, allele *S3* is recessive to all other alleles which are co-dominant with each other (*S3* < *S1* = *S2* = *S4*). **(B)** PCR-amplification with allele specific *PdPIRK* primers using gDNA for all ten individuals reveals that *PdPIRK* genotype corresponds perfectly with inferred *S*-locus genotype.

To determine genotypic variation at *PdPIRK*, we generated long-read transcriptomes of stigma/style tissue for one individual of each cross-compatibility group and recover two alleles of *PdPIRK* in each transcriptome. As expected, these four alleles display high polymorphism (Nucleotide Pairwise Identity = 74.3%). Leveraging this exceptional variation, we developed a presence/absence PCR-genotyping assay using primers designed to specifically and uniquely amplify each allele (Supplemental Table S3). For all ten individuals in our diallel cross, genotypic variation at *PdPIRK* perfectly predicts crossing success among these individuals (Figure 3B). We refer to these alleles of *PdPIRK* as *S1*, *S2*, *S3*, and *S4* enumerated by the order in which alleles were identified and sequenced.

### *PdPIRK* controls compatibility in an allele-specific pattern

We functionally validate that *PdPIRK* causes self-pollen recognition using virus-induced gene silencing (VIGS) to knockdown its expression^28^. *Phlox* displays developmental variation in the SI response which we exploited to force self-fertilization of one *S2S4* plant and produce a family of individuals with possible genotypes *S2S2, S2S4,* and *S4S4*. We used our long-read transcriptomes to design a silencing vector specifically targeting allele *S4* and infected *S4S4* homozygous individuals (n=5). VIGS is heterogenous such that not every flower on an infected plant will experience silencing. We therefore co-silence an essential gene in the anthocyanin pigment biosynthesis pathway, dihydroflavonol 4-reductase (*DFR*), as a marker to indicate silencing in floral tissue (Figure 4A&B; Supplemental Figure S4)^29^. As confirmed by qPCR, stigma/styles from visibly white flowers have reduced *S4* expression compared to pistils from pigmented flowers (% expression of pigmented flowers, mean ± SE = 38.4 ± 13.5) (Figure 4C). Following self-pollination, pigmented/unsilenced flowers set a single fruit in all crosses across all replicate plants, compared to white/silenced flowers that set fruit in 60% (SE = 10.75%) of self-pollinations (t_fruit set≠0_ (4) = 5.58, p = 0.0051) (Figure 4D). This observed increase in self-compatibility is mediated by a reduced SI response at the pollen-pistil interface as demonstrated by quantifying the number of pollen tubes germinated on silenced vs. unsilenced stigmas (z = - 4.82, p < 0.001) (Figures 4A&E). We replicate this result using a silencing vector targeting allele *S1* in *S1S1* plants (n=4). As confirmed by qPCR, stigma/styles from visibly white flowers have reduced *S1* expression compared to pistils from pigmented flowers (% expression of pigmented flowers, mean ± SE = 27.3 ± 9.86) (Figure 4C). We again observe a significant increase in fruit set and pollen tube germination following self-pollination in silenced vs. unsilenced flowers (fruit set mean ± SE = 56.2 ± 12; t_fruit set≠0_ (3) = 4.70, p = 0.018; pollen tube z = -4.21, p < 0.001) (Figures 4A,F&G). Control experiments silencing *DFR* alone reveal no effect of our experimental manipulation protocol on self-fruit set (t_fruit set≠0_ (5) = 1, p = 0.3632) (Supplemental Figure S3). We therefore conclude that *PdPIRK* is a necessary component of functional self-incompatibility.

**Figure 4:**
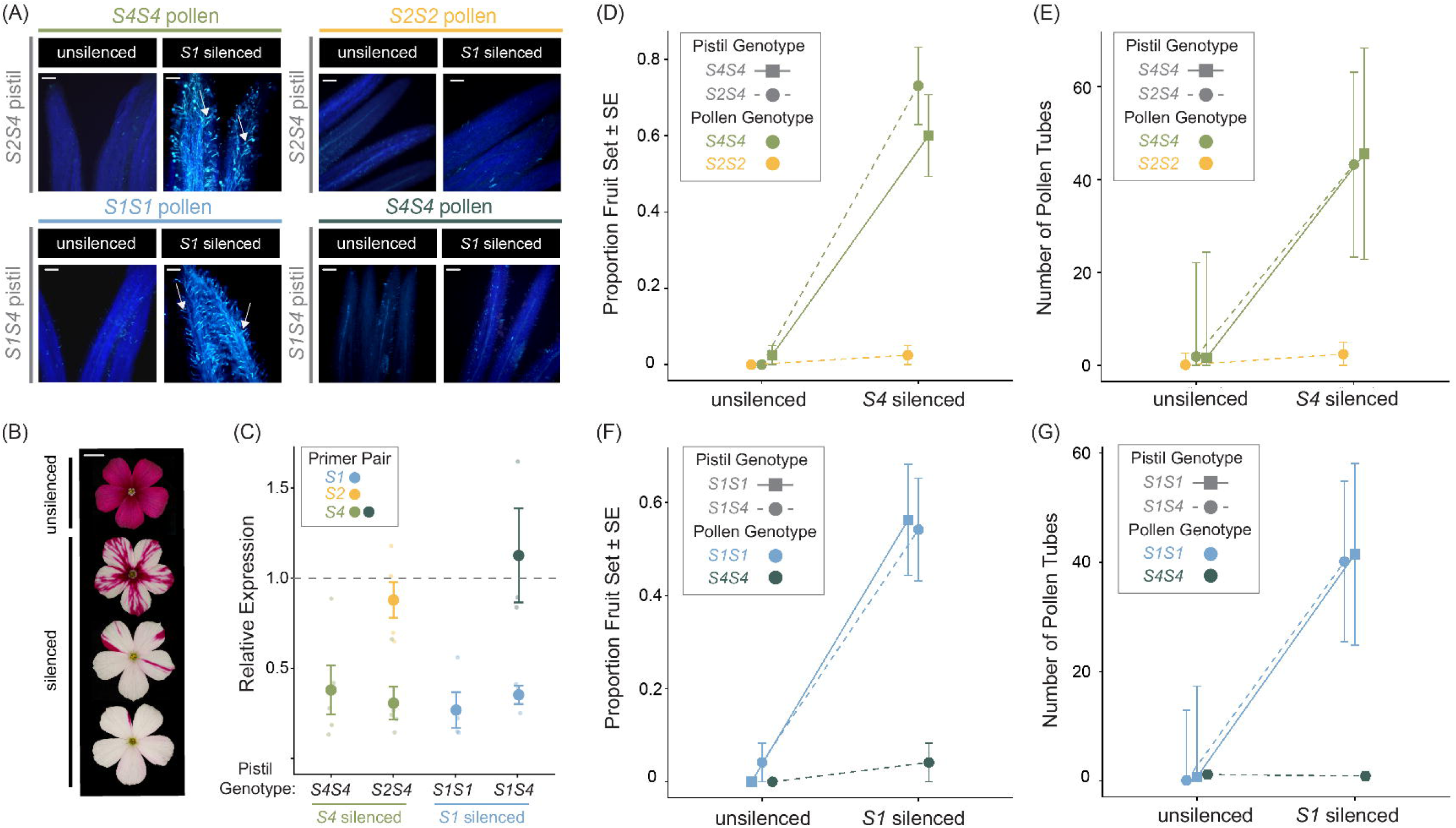
Silencing *PdPIRK* affects compatibility in a genotype-dependent pattern. **(A)** Representative images of stigmas following pollination for each treatment type. White arrows indicate pollen tubes. Pollen tubes do not germinate on stigmas from pigmented/unsilenced flowers for any cross type. For stigmas from *S4* silenced flowers of heterozygous *S2S4* plants, pollen tubes germinate if pollen is *S4S4* but not *S2S2.* For stigmas from *S1* silenced flowers of heterozygous *S1S4* plants, pollen tubes germinate if pollen is *S1S1* but not *S4S4* (scale bar = 100um). **(B)** Unsilenced flowers are fully pigmented while silenced flowers are white to varied degrees (scale bar = 1cm). **(C)** Percent expression of *PdPIRK* in stigma/style tissue from white flowers compared to pigmented flowers as quantified by qPCR. Mean ± SE is plotted along with sample data. Points are colored based on the *PdPIRK* allele amplified, with *S1* shown in blue, *S2* in yellow and *S4* in greens. **(D)** Silencing *PdPIRK* allele *S4* specifically causes an increase in fruit set and **(E)** pollen tube number depending on pollen genotype. **(F)** Silencing *PdPIRK* allele *S1* specifically causes an increase in fruit set and **(G)** pollen tube number depending on pollen genotype. Means ± standard errors are plotted for fruit set and model predictions with 95% confidence intervals are plotted for pollen tube number. Homozygous pistil genotypes are shown as squares with solid lines and heterozygous pistil genotypes are shown as circles with dashed lines. Colors correspond to pollen genotype.

We demonstrate that *PdPIRK* activity is allele-specific, confirming its role in recognizing self-pollen identity per se, as opposed to operating elsewhere in the SI pathway. To functionally validate that *PdPIRK* controls recognition, we infect *S2S4* heterozygous plants (n=5) with a silencing vector uniquely targeting allele *S4.* Confirmed by qPCR, this treatment results in reduced *S4* expression in white versus pigmented flowers while *S2* expression is much less affected (% expression of pigmented flowers, mean ± SE = 31.1 ± 9.1 for *S4*; 88.1 ± 9.84 for *S2*) (Figure 4C). As predicted, silencing allele *S4* causes an increase in compatibility with *S4S4* homozygous pollen (mean ± SE = 73.1 ± 10.13; t_fruit set≠0_ (4) = 7.215, p = 0.002). In contrast, *S2S2* pollen is consistently recognized as self and rejected by pistils from silenced flowers (mean ± SE = 2.5 ± 2.5; t_fruit set≠0_ (4) = 1, p = 0.374) (Figure 4D). Pollen tube data similarly reveal such allele-specific activity (*S4S4* pollen z = -4.12, p < 0.001; *S2S2* pollen z = -1.67, p = 0.095) (Figures 4A&E). We replicate this result by uniquely silencing allele *S1* in *S1S4* plants (% expression of pigmented flowers, mean ± SE = 35.7 ± 5.1 for *S1*; 112.8 ± 26.0 for *S4*). As predicted, we observe a significant increase in compatibility with *S1S1* homozygous pollen and no increase in compatibility with *S4S4* homozygous pollen (*S1S1* pollen mean ± SE = 54.2 ± 11.0; t_fruit set≠0_ (2) = 4.24, p = 0.032; *S4S4* pollen mean ± SE = 4.2 ± 4.2; t_fruit set≠0_ (2) = 1, p = 0.423) (Figure 4F). Pollen tube data similarly reveal such allele-specific activity (*S1S1* pollen z = -3.61, p < 0.001; *S4S4* pollen z = 0.495, p = 0.62) (Figures 4A&G). We therefore conclude that *PdPIRK* controls self-pollen recognition in *Phlox*.

### *PdPIRK* encodes a lectin receptor-like kinase

*PdPIRK* is annotated as a receptor-like kinase (RLK), a large and widespread family of transmembrane proteins in flowering plants^30^. Amino acid sequence analysis suggests that *PdPIRK* is a member of the G-type subclass of lectin receptor-like kinases (LecRLKs) in particular, a group of cell surface receptors known for their ability to perceive and transduce signals from the environment^31, 32^. The G-type subclass is marked by an extracellular region containing a bulb-type lectin, *S*-locus glycoprotein, and Plasminogen/Apple/Nematode (PAN) domain^30, 33^. We reconstruct evolutionary relationships among protein sequences containing these domains (G-type LecRLKs) and related sequences containing legume lectin domains (L-type LecRLKs)^34^. Using sequences from ten plant genomes, we find that *PdPIRK* is nested within the G-type LecRLK clade, corroborating its identity as a member of this subclass (Supplemental Figure S6; Supplemental Table S6). Notably, the gene responsible for self-pollen recognition in the distantly related Brassicaceae (*SRK*) is also a G-type LecRLK. (Figure 1A; Supplemental Figures S5&S6; Supplemental Table S6)^35, 36^.

## Discussion

Here, we reveal the identity of a previously uncharacterized *S*-locus by presenting multiple, independent lines of evidence that *P. drummondii* Pistil Identity Receptor Kinase (*PdPIRK)* causes self-pollen recognition in *Phlox*. *PdPIRK* is located within the genomic region associated with the *S*-locus in *Phlox* and is the only gene in this region to match two key predictions for how *S*-loci function and evolve^18^. First, we find a pattern of high expression localized to stigma/style tissue, the site where self-pollen recognition occurs. Second, reconstructing evolutionary relationships among alleles both within and between species demonstrates exceptional polymorphism, a classic signature of negative-frequency dependent selection. Moreover, we find a perfect association between *PdPIRK* genotype and compatibility phenotype in a diallel cross among individuals sharing *S*-alleles. Finally, we functionally validate *PdPIRK* by knocking down its expression which increases compatibility in an allele-specific pattern confirming its role in recognizing pollen identity.

Homomorphic *S*-loci have evolved independently a minimum of 14 times throughout flowering plant diversification^7^. Our identification of *PdPIRK* represents an apparent case of genetic convergence among these *S*-loci; self-pollen recognition in *Brassica* is also caused by a G-type LecRLK^11, 36^. Among the five previously characterized homomorphic *S*-loci, each achieves self-pollen recognition via unique molecular pathways, demonstrating that the repeated evolution of SI can and does proceed through nonparallel genetic routes^12–14, 36^. Our findings suggest that this lack of convergence is not a persistent feature of homomorphic *S*-locus evolution. Recent identification of novel *S*-loci in heteromorphic SI systems similarly reveals such molecular parallelism. For example, the *S*-loci of *Primula, Turnera* and likely *Linum* contain brassinosteroid inactivating enzymes which enforce outcrossing by generating dimorphic floral forms^37–39^. Collectively, the repeated co-option of members of the same gene families to function in SI suggests the presence of constraints or paths of least resistance in the kinds of genes that can evolve to cause self-pollen recognition.

The self-incompatibility response in *Brassica* and *Phlox* share additional physiological characteristics. Both taxa produce dry stigmas, reject self-pollen prior to stigma binding, and encode pollen identity sporophytically^18, 40, 41^. These observations motivate future research exploring the extent of convergence between the two SI mechanisms. Identification of interacting molecules in the *Phlox* SI pathway, including the pollen-expressed ligand that binds *PdPIRK*, will clarify whether similar genes operate throughout the pollen rejection response in both systems. Several species with currently unknown *S*-loci possess similar reproductive features (e.g., species within the Asteraceae) and our findings suggest that G-type LecRLKs may be promising initial gene candidates^42–44^.

Molecular mechanisms of self/non-self-discrimination are essential to diverse biological processes. Characterizing the genetic basis of self-recognition contributes insight into the function and evolution of these systems and is a foundational step towards future modifications or manipulations. Identifying the genetic control of self-pollen recognition has wide-spread applications for agriculture and conservation. For example, crop yields, breeding program success, and the likelihood of domestication are all influenced by a plant’s ability to self-fertilize^45^. For threatened and endangered species, knowledge of the genetic basis of SI can be applied during management to alleviate mate limitation and increase the likelihood of species persistence^46–48^.

Independent evolutions of SI provide natural replication of evolutionary processes which can be leveraged to reveal patterns in how self-recognition evolves. Given the limited number of known *S*-loci, it appeared that the common phenotype of homomorphic SI represented a constellation of nonparallel genetic mechanisms; our identification of *PdPIRK* suggests that convergence in *S*-locus genes may be more common than previously seemed^19^. Our study demonstrates how using specific predictions for *S*-locus evolution can narrow candidate genes and pinpoint the underlying genetic control of self-pollen recognition^22^. We outline a strategy for identifying additional *S*-loci across diverse and non-model taxa which will elucidate the varied mechanisms plants use to achieve self-recognition while clarifying the extent of molecular convergence and constraint in the evolution of SI.

## Materials and Methods

### Plant collection and care

We collected seeds from naturally occurring *P. drummondii* in May of 2019 which served as parents for the families used in this study (Supplemental Figure S1). All plants were grown under standardized conditions at the Arnold Arboretum of Harvard University. We germinated plants by first incubating seeds in 500ppm gibberellic acid in water for two days and subsequently cold stratified planted seeds at 4°C for ten days. Plants were then transferred to 4-inch pots with Pro-Mix HP Myccorhizae potting media and maintained in growth chambers kept at a daytime temperature of 23^0^C and a nighttime temperature of 18^0^C, with 16 hours of supplemented light.

### Analyses of tissue-specific expression

We accessed mRNA sequence data of five tissue types (stigma/style, anther, pollen, leaf, stem) from Burgin et al. (2025)^18^. We aligned reads to the *P. drummondii* genome assembly (NCBI BioProject: PRJNA1219593, unpublished) using a 2-pass alignment with STAR v2.7.11b^49^. Abundance calls were generated using RSEM v1.3.0 and used as input for downstream analyses in R v.4.2.3^50^. Read counts were normalized across samples using the TMM method. Genes with restricted expression in stigma/style tissue were identified by characterizing differentially expressed genes pairwise between stigma/style samples and each other tissue type independently in edgeR^51, 52^. The quasi-likelihood *F*-test was used to identify differentially expressed genes using the glmQLFTest function. We adjusted p-values via the Benjamini-Hochberg Procedure. Genes meeting the criteria of 1) greater than 5-fold change and 2) a significance threshold of p < 0.01 in all pairwise tissue comparisons were considered restricted in expression to the stigma/style.

The genomic region containing the *S*-locus in *P. drummondii* contains 727 genes. Based on normalized expression averaged across biological replicates (n=6), the most highly expressed gene in stigma/style tissue within this region is *Pd52568* which we name *PdPIRK*. This gene is also restricted in expression to stigma/style tissue, and we confirm this pattern using quantitative real time PCR (qPCR) of leaf, corolla, and stigma/style tissue from three biological replicates of *S4S4* homozygous plants (see below for genotyping methods). Tissue was collected under RNase-free conditions and flash-frozen on liquid nitrogen. A single leaf and corolla were collected from each plant. To collect stigma/style tissue, we used fine-tip forceps to dissect out the pistil from the flower and then remove the ovary. We pooled stigma/style tissue from eighteen flowers of each plant collected on the first or second day the flower opened. All tissue was stored at -80°C until RNA extraction.

We finely ground tissue on liquid nitrogen with a sterile pestle and extracted total RNA using the Spectrum Total RNA Kit (Sigma-Aldrich, St. Louis, Missouri, USA). We removed DNA by performing an on-column digestion using the DNase I Digestion Set (Millipore Sigma, Burlington, MA, USA). Yield and purity were assessed by Nanodrop before synthesizing cDNA via the NEB ProtoScript First Strand cDNA Synthesis Kit (New England Biolabs, Ipswitch, MA, USA).

Each cDNA sample was amplified and quantified using Brilliant SYBR Green qPCR Mastermix (Agilent, Santa Clara, CA, USA) with primers targeting *PdPIRK* allele *S4* and the housekeeping gene, *PdEf1α*. All samples were amplified for 40 cycles with an annealing temperature of 60°C. Critical threshold (Ct) cycle values were obtained using the Stratagene Mc3005P instrument (Agilent, Santa Clara, CA, USA). We performed two or three technical replicates for each amplification and averaged Ct values across these replicates. The difference between the average critical threshold values for *PdEf1α* and *PdPIRK* was calculated (ΔCt) and linearized (2^-ΔCt^). To determine if *PdPIRK* expression varied across tissue types, we performed a two-way analysis of variance (ANOVA) and evaluated significant differences with post-hoc Tukey HSD tests.

### Phylogenetic analyses of polymorphism

Because we have an *a priori* expectation that the gene responsible for self-pollen recognition will be among the most highly expressed in stigma/style tissue genome-wide, we focus analyses of polymorphism to the top ∼10% of genes most highly expressed within the *S*-locus associated region (n=73). We used our stigma/style transcriptomes to generate twelve phased alleles for each of these 73 genes. To call variants, we used the GATK4 Joint Genotyping best practices workflow for RNAseq data^53^. We filtered variants to include only single-nucleotide polymorphisms with GQ > 10 and a minimum read depth of 5 using SAMtools v1.17 and phased variant calls using WhatsHap v2.3^54, 55^. We reconstructed alleles by extracting predicted coding sequences from the *P. drummondii* reference genome assembly and using SAMtools to merge with the phased variant calls.

Nucleotide sequences were converted to predicted amino acid sequences and aligned using the ClustalW algorithm in Geneious v11.0.24. Phylogenetic relationships among alleles for each gene were estimated using random accelerated maximum likelihood (RAxML) with the LG+G+F model of amino acid replacement ^56^. For each gene, the total branch length (substitutions per site) was quantified for the best scoring tree. To formally test for statistical outliers, we implemented Rosner’s Test for Outliers using the *EnvStats* package in R v.4.2.3^57^. To ensure that variation in sequence coverage between genes does not bias our ability to identify variants and ultimately branch lengths, we tested for a correlation between average gene expression in stigma/style tissue and total gene tree branch length and found no correlation when *PdPIRK* is removed from the dataset (Supplemental Figure S2; adjusted R-squared = -0.0078, p = 0.503).

To determine whether *PdPIRK* displays a pattern of trans-specific polymorphism, we curated long-read sequencing from (NCBI BioProject: PRJNA1219593, unpublished) of four closely related *Phlox* species: *P. drummondii* (n=3), *P. roemeriana* (n=3), *P. cuspidata* (n=4), and the outgroup *P. pilosa* (n=1). High molecular weight DNA was extracted and sequenced via Oxford Nanopore Technologies, and raw reads processed as in Garner et al.^58^. Reads were aligned to the *P. drummondii* reference assembly using minimap2 v2.28-r1209^59^. We called SNPs using mpileup and call algorithms in bcftools suite v1.17 and removed SNPs with QUAL ≤ 20 and/or read depth ≤ 5^54^. We applied called variants to the predicted coding sequences of the *P. drummondii* reference genome using the consensus command in bcftools for three genes within the *S*-locus associated region: *PdPIRK*/*Pd52568*, *Pd51610*, and *Pd51965*. We aligned nucleotide sequences using the ClustalW algorithm in Geneious v11.0.24. Phylogenetic relationships among alleles for each gene were estimated using random accelerated maximum likelihood (RAxML) with the LG+G+F model of nucleotide replacement^56^.

### Associating *PdPIRK* genotype with cross-compatibility phenotype

To test whether genotypic variation at *PdPIRK* predicts cross-compatibility phenotype, we began by crossing two individuals to generate n=10 full sibling offspring (given IDs A0 – A9) with four possible *S*-locus genotypes. For clarity, we refer to parent 1’s alleles as *S1* and *S2* and parent 2’s alleles as *S3* and *S4*. We performed a full diallel cross among all ten offspring to infer *S*-locus genotype based on cross-compatibility. Specifically, we expect that crosses between individuals sharing *S*-alleles will be incompatible and produce no fruit while crosses between individuals with different *S*-alleles will be compatible and produce fruit.

Crosses were performed by pollinating flowers 1-2 days following floral opening using forceps to transfer pollen onto stigmas. Each flower can produce a single fruit, and we pollinated six flowers per cross for a total of six possible fruits. We quantified fruit set a minimum of 1-week post-pollination to ensure fruit had time to develop. We previously showed that variants conferring self-compatibility segregate within *P. drummondii* which can confound patterns of cross-compatibility^18^. Therefore, we screened plants for complete SI by performing eight self-pollinations per plant and found that all individuals set zero self-fruits.

After organizing individuals into groups with similar crossing behavior, we find four cross-compatibility groups consistent with four possible *S*-locus genotypes. As is known for other sporophytic SI systems, we find dominance interactions among alleles. In this cross, allele *S3* is recessive to all other alleles which are co-dominant with each other (*S3* < *S1* = *S2* = *S4*). To test whether *PdPIRK* genotype correlates with inferred *S*-locus genotype based on crossing phenotype, we performed full-length transcript RNA sequencing on stigma/style tissue for one individual of each cross-compatibility group. Based on these sequences, we designed primers to specifically amplify each of the four *PdPIRK* alleles recovered. We used these primers to PCR-amplify genomic DNA (gDNA) from each individual and determine *PdPIRK* genotype based on presence/absence of amplified DNA (see section “PacBio Iso-Seq of stigma and style tissue” for details).

### PacBio Iso-Seq of stigma and style tissue

We collected stigma/style tissue using fine-tip forceps to dissect out the pistil from the flower and remove the ovary. We pooled tissue from eighteen flowers for one plant of each inferred *S*-genotype (plant IDs: A5, A6, A7, A9). Pistils were collected from flowers on the first or second day the flower opened, and tissue was stored at -80°C until RNA extraction. We finely ground tissue on liquid nitrogen with a sterile pestle and extracted total RNA using the Spectrum Total RNA Kit (Sigma-Aldrich, St. Louis, Missouri, USA). We removed DNA by performing an on-column digestion using the DNase I Digestion Set (Millipore Sigma, Burlington, MA, USA). Yield and purity were assessed by Nanodrop, and samples were screened for RNA degradation by TapeStation (Agilent, Santa Clara, California, USA).

Full-length transcript sequencing was performed by Novogene using the PacBio SMRT sequencing platform (PacBio, Menlo Park, California, USA). We used RNA-Bloom2 to assemble HiFi reads into de novo transcriptomes which were queried against the *PdPIRK/Pd52568* sequence from the *P. drummondii* genome using BLASTn^60, 61^. We predict each plant to express two unique *PdPIRK* alleles and our short-read RNAseq demonstrates that its expression is extremely high in the stigma/style. Therefore, we extracted sequences of the two hits with highest read coverage. In this process, we recovered sequences from individuals as predicted by cross-compatibility (e.g., individual A7 shared one sequence with A9 and no sequences with A6). For each allele, we designed 25bp primers that uniquely bind to one allele but not the other three. A modified CTAB extraction was used to extract gDNA from leaf tissue for all ten individuals in our diallel cross. Each sample was amplified for 28 cycles with an annealing temperature of 65.5°C for primer pairs *S2* and *S3* and 66.5°C for primer pairs *S1* and *S4*. Presence/absence of DNA amplification was assessed by electrophoresis (Fig. 2C).

### Virus-induced gene silencing (VIGS) of *PdPIRK*

VIGS constructs were initially acquired from the Arabidopsis Biological Resource Center (pTRV1 accession CD3-1039; pTRV2-MCS accession CD3-1040) ^28^. Based on the *Phlox drummondii* genome assembly, we identified a 300bp sequence unique to the coding region of dihydroflavonol 4-reductase (*DFR*). We used Twist Biosciences (South San Francisco, California, USA) to synthesize this sequence flanked by BamHI and EcoRI digestion sites. Synthesized fragments were cloned into the pTRV2-MCS plasmid by restriction cloning and used for the *DFR* only control experiment. To functionally validate the effect of *PdPIRK* on self-incompatibility, we used our long-read stigma/style transcriptomes to design a 225bp sequence uniquely targeting the *S4* and *S1* alleles separately. We used Twist Biosciences to synthesize a fragment containing either sequence flanked by XmaI and MspI digestion site, and restriction cloning to insert fragments 3’ of the *DFR* target sequence into the pTRV2-DFR plasmid.

We electroporated pTRV2-DFR, pTRV2-DFR-PdPIRK(S4), pTRV2-DFR-PdPIRK(S1), and pTRV1 constructs into *Agrobacterium tumefasciences* (GV3101) and positively selected transformants by growing colonies overnight at 28°C on YEB agar plates containing kanamycin, gentamycin, and rifampicin antibiotics. Colonies carrying the correct construct were initially verified using PCR and confirmed with direct sequencing by Eton Bioscience (San Diego, California, USA). We used single colonies to inoculate 3mL overnight YEB cultures grown at 28°C in the presence of kanamycin and gentamycin antibiotics which was subsequently used to inoculate a 50mL overnight culture incubated under the same conditions. To prepare bacteria for inoculation, we pelleted by centrifugation and resuspended cells in freshly prepared infiltration buffer (10mM MES, 200uM acetosyringone, 10mM MgCl_2_). All cultures were normalized to OD_600_=10 and pTRV1 cells were separately mixed in a 1:1 volume with pTRV2-DFR, pTRV2-DFR-PdPIRK(S4) or pTRV2-DFR-PdPIRK(S1) cells. These mixtures were incubated at room temperature with nutation for 2-3 hours before transforming plants.

Plants were grown for 1-month post-germination under the standardized conditions described above. To control for plant fertility, we screened all individuals for outcross fruit set and pollen viability. Individuals setting fewer than five fruits following outcross pollination using pollen from an unrelated individual (out of eight total flowers crossed) and/or less than 90% pollen viability were removed from the experiment. Pollen viability was quantified using a modified Alexander’s Stain protocol^62^. Briefly, we collected pollen from three flowers per plant and stored samples overnight at 4°C in 70% ethanol. Alexander Stain was added to achieve a final concentration of 0.01% Malachite Green, 0.05% Acid Fuschin, 0.005% Orange G, 4% glacial acetic acid, 25% glycerol, and 10% ethanol. Pollen was incubated in the stain for ∼6 hours at room temperature. We placed a 50uL volume onto a glass slide and heated over a Bunsen burner until liquid evaporated. Pollen was visualized in the brightfield and rhodamine (544nm) channels at 5x magnification using an Axio Imager.A2 (Zeiss, Jena, Germany) microscope with ZEN 2.3 pro software. We automated quantification of the proportion of viable pollen grains in ImageJ 1.53k and confirmed all counts manually ^63^. Pollen grains were considered viable if they stained purple and fluoresced in the Rhodamine channel. Because our previous work demonstrates variation in the SI response segregating within *P. drummondii*, we further screened plants for complete SI. Individuals setting any fruit following self-pollination (out of eight total flowers crossed) were removed from the experiment.

To encourage lateral meristem initiation throughout the plant, branches were pruned three days prior to transformation. We used 28-guage needles to inject the pTRV1 + pTRV2-DFR, pTRV2-DFR-PdPIRK(S1) or pTRV1 + pTRV2-DFR-PdPIRK(S4) inoculum into all visible axillary meristems. A total of ten plants were treated with pTRV1 + pTRV2-DFR, ten with pTRV1 + pTRV2-DFR-PdPIRK(S4) (n=5 *S2S4* plants and n=5 *S4S4* plants), and ten with pTRV1 + pTRV2-DFR-PdPIRK(S1) (n=5 *S1S4* plants and n=5 *S1S1* plants). Twenty-six of the thirty treated plants began producing white flowers 2-5 weeks after transformation. We considered a flower white if approximately 25% or more of the corolla was visibly unpigmented. To confirm that white flowers indicate successful *PdPIRK* silencing in pTRV2-DFR-PdPIRK(S4) and pTRV2-DFR-PdPIRK(S1) treated plants, we collected stigma/style tissue for each individual by pooling 18 open flowers of either floral color type (white or pigmented). Total RNA was extracted and converted to cDNA and we used quantitative real time PCR to quantify expression of *PdPIRK* allele *S4* or *S1* as described above. For heterozygous plants, we additionally quantified expression of *PdPIRK* allele *S2* or *S4* to confirm allele specific silencing. Expression of the housekeeping gene, *PdEF1a*, was used to normalize *PdPIRK* expression in each sample. Standard curves were run for all primer pairs to confirm efficiency. The relative expression of *PdPIRK* in white versus pigmented flowers was calculated for each plant using the ΔΔCt method^64^.

### Quantifying the effect of silencing *PdPIRK* on self-incompatibility

We quantified the effect of silencing *PdPIRK* allele *S4* on SI by performing a series of controlled pollinations assessing both fruit set and pollen tube germination. Cross types in this experiment include *S4S4* homozygous plants receiving self-pollen, *S2S4* heterozygous plants receiving pollen from an *S2S2* homozygous plant, and *S2S4* heterozygous plants receiving pollen from an *S4S4* homozygous plants. We repeated this experiment with silencing of *PdPIRK* allele *S1* and performed the following cross types: *S1S1* homozygous plants receiving self-pollen, *S1S4* heterozygous plants receiving pollen from an *S1S1* homozygous plant, and *S21S4* heterozygous plants receiving pollen from an *S4S4* homozygous plants.

To assess fruit set, eight pigmented/unsilenced and eight white/silenced flowers per individual were pollinated for each cross type. Because all plants included in the experiment were initially screened for complete SI, removing anthers pre-dehiscence to prevent incidental self-pollination was unnecessary. Flowers were pollinated 1-2 days following floral opening by using forceps to transfer pollen onto stigmas. We quantified fruit set a minimum of 1-week post-pollination to ensure fruit had time to develop. Each flower can produce a single fruit which we recorded as binary outcome data (1 for fruit, 0 for no fruit). For several of our treatments, zero unsilenced flowers set fruit in our experiment which rendered model-based statistical approaches infeasible due to the lack of variance. We therefore implemented a Student’s t-Test in R v.4.2.3 to determine if fruit set in silenced treatments was statistically different from zero for each cross type separately.

To assess pollen tube germination, four pigmented/unsilenced and four white/silenced flowers per individual were pollinated for each cross type. We removed calyces with attached pistils from plants 24 hours after pollination and submerged tissue in formalin acetic acid (FAA) for an additional 24 hours at room temperature. Tissue was transferred to 70% ethanol for long-term storage at 4°C. Following a minimum of 1 month in 70% ethanol, tissue was incubated in 5M sodium hydroxide overnight at room temperature. Pistils were then dissected out of calyces, stained in 0.1% aniline blue in .1N K_3_PO_4_ for 2 hours, and mounted on glass slides in 50% glycerol. We visualized pollinated pistils in the DAPI channel (358nm) at 10x magnification using an Axio Imager.A2 (Zeiss, Jena, Germany) microscope with ZEN 2.3 pro software, and manually quantified the number of pollen tubes present. We implemented a generalized linear mixed effects model in R v.4.2.3 using the R package *lme4* v. 1.1-34 ^65^. Our model included flower color (pigmented/unsilenced vs. white/silenced) as a fixed effect and plant ID as a random effect.

### Characterizing homology with G-type subclass LecRLKs

We generated amino acid sequences of *PdPIRK* alleles identified by IsoSeq and annotated predicted protein motifs using DeepTMHMM and SIB Swiss Motif Scan^66, 67^. We queried the *P. drummondii* genome against the *PdPIRK* using BLASTn to identify an additional lectin domain containing receptor-like kinase sequence, *Pd41729*, which is annotated outside of the *S-*associated region^60, 61^. Lectin domain containing receptor-like kinase sequences from diverse species were identified using InterPro (G-type LecRLKs: IPR024171; L-type LecRLKs: IPR050528) from ten plant genomes (*Oryza sativa, Zea mays, Arabidopsis thaliana, Latuca sativa, Capsella grandiflora, Brassica rapa, Brassica napus, Brassica oleraceae, Aquilegia coerulea,* and *Vitis vinifera*). We included two additional accessions of known *SRK* sequences from NCBI (AB054720.1, Z30211.1) (Supplemental Table S6). Sequences were aligned using the ClustalW algorithm in Geneious v11.0.24. Phylogenetic relationships among alleles for each gene were estimated using random accelerated maximum likelihood (RAxML) with the LG+G+F model of amino acid replacement^56^.

## Data availability

Raw sequencing reads for tissue-specific RNA sequencing can be accessed via the NCBI sequence read archive (SUB15344310). The *Phlox drummondii* genome assembly and annotation are hosted by Zenodo and accessible at (doi: 10.5281/zenodo.15498773). Long-read DNA sequences used to construct gene trees for the four *Phlox* species are available via NCBI (BioProject: PRJNA1219593). Long-read RNA sequences of stigma/style transcriptomes are deposited in the NCBI sequence read archive (SUB15389546) and assembled transcriptomes are available via the Dryad Digital Repository (doi: 10.5061/dryad.zgmsbccr3). All other phenotyping data including fruit set, pollen tube germination, floral scans for flowers used in silencing experiments, qPCR data, and all sequence alignments used to construct gene trees are available via the Dryad Digital Repository (doi: 10.5061/dryad.zgmsbccr3). All code required for the analyses presented here are available at: https://github.com/graceburgin/phlox_SI. Any additional information necessary to reanalyze data reported in this paper is available from corresponding authors upon request.

## Authors’ contributions

G.A.B. and R.H. conceived of the study. N.L. designed and performed the diallel crossing experiment. G.A.B. designed and performed the remaining experiments, and R.H. and N.L. participated in data collection efforts. G.A.B. performed all data analyses and wrote the manuscript together with R.H. All authors contributed to editing and revising the manuscript.

## Acknowledgements

We thank Andrea Berardi, Patrick McKenzie, Christina Steinecke, Boris Igić, Elena Kramer and Felix Wu for guidance and comments on the study. Essential plant care was provided by the Arnold Arboretum Greenhouse Staff, and the authors especially thank Megan Ardolino, Mike Barrett, Aberdeen Bird, and Maya Levine.

