## Supplemental Figures for "A lectin receptor-like kinase controls self-pollen recognition in *Phlox* (Polemoniaceae)"

| Individual ID | Population ID | Year Sampled | Latitude | Longitude |
|---|---|---|---|---|
| 82 | 679 | 2019 | 29.367483 | -97.566617 |
| 6 | 667 | 2019 | 29.528817 | -96.441233 |
| 21 | 664 | 2019 | 30.116 | -97.365 |
| 234 | 656a | 2019 | 30.0852333 | -96.916167 |
| 88 | 676 | 2019 | 29.4689024 | -97.873746 |
| 226 | 679 | 2019 | 29.367483 | -97.566617 |
| 154 | 670 | 2019 | 29.4174167 | -97.6808 |

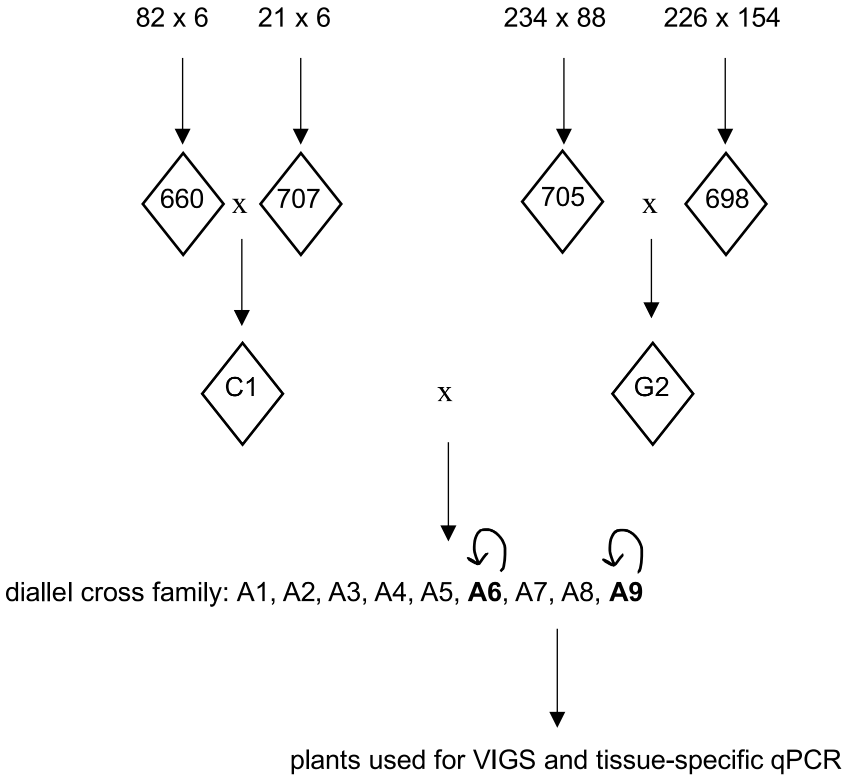


**Figure S1.** Sampling and line information for plants used in this study. Individual IDs are given along with GPS coordinates for original plants sampled from natural populations of *Phlox drummondii*. Crosses are indicated by an x with the pistil parent listed first and arrows leading to offspring. The circular arrows indicate forced selfing of individuals A6 and A9 to produce all offspring used for VIGS and tissue-specific qPCR experiments.

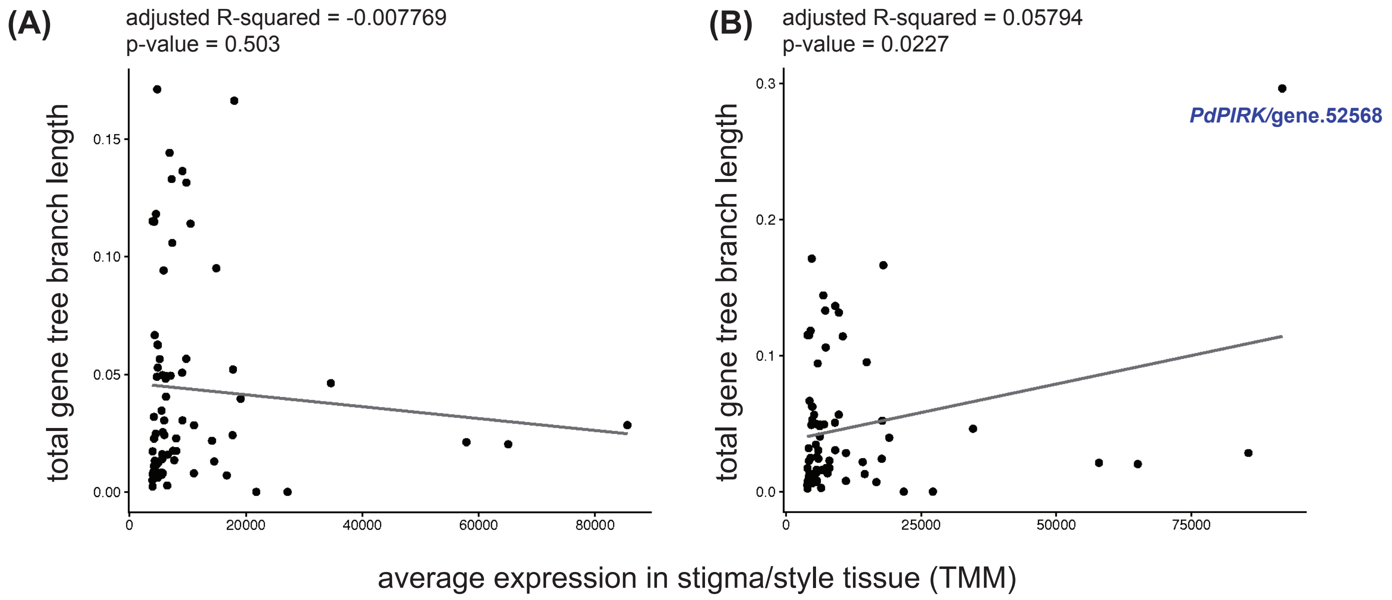


**Figure S2.** The relationship between average expression in stigma/style tissue and total branch length for **(A)** the n=73 most highly expressed genes in stigma/style tissue annotated to the S-locus associated region with *PdPIRK* removed from the dataset and **(B)** including *PdPIRK* in the dataset.

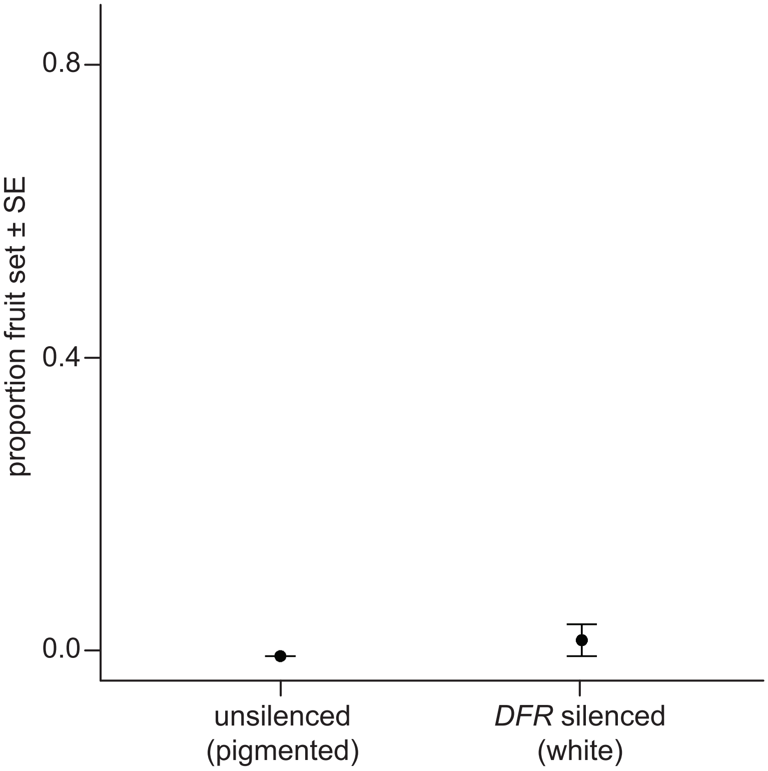


**Figure S3.** Silencing *DFR* alone using VIGS does not cause an increase in fruit set following self-pollination ($t_{fruit\ set \neq 0}$ (5) = 1, p = 0.3632). Means are plotted as points with standard errors.

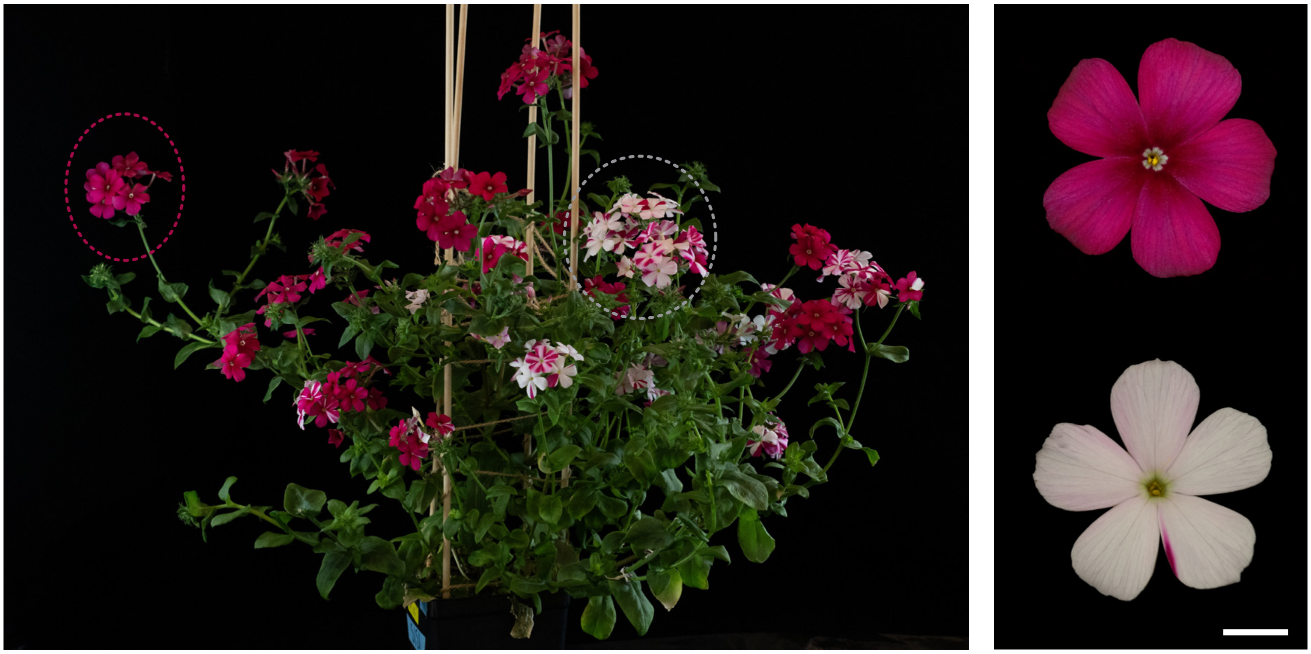

**Figure S4.** Floral phenotypes of *PdPIRK*(*S4)-DFR* VIGS knock-down in *P. drummondii*. Pigmented flowers (circled in pink) and white flowers (circled in gray) are produced on branches of the same plant (scale bar = 1cm).

(A)

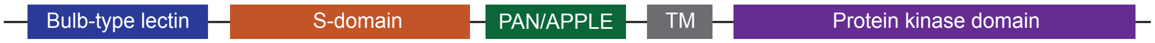


(B)

MGARGRKNTMHDFRKSLIFFILLLLIEFSSGRDTLSNGETLNDKDGDSIISAKKTFTLAFFSPGMSPRRYLAISYYSLPENKVVW

VANRDKPVNDSSAVLSFHTPNNLVMYGSDLKVPIWSTSFLGPGKNYSARLLDTGNLVVFEEGNRGFIAWQSFDHPTDTWLP

EMKFGVNRKTGLSWSLRSWKSFDDPGTGEYSLRMDGENHELSGLTLYKGLHRIWRSGPLTGGGQFMADMSRDSSLNLTYE

SNENEVYLMFTVHNLMNSIMVVNYLGNAVRRTMLPGAPRWRGVWKAPEGLCDHYGMCGPNRNCDAGSTAPLINCECLP

GSKIKSQREFDAGNMSGGCVRNRDRNDCWNGEGFTKLSGTEMPDTASARVDKMRGSRECEEECLRDCNCTAYTNVPKMG

NACVMWYGDLMDTKQYGEGGLDLFVRDVNGTRNMTQQQHYSSNKKRRVAAVVTSVGVVAFILVFILCWLTYKRKGGLRSR

AEQRNGNLLNGPDNSTDQIVGRQTSGDLPFFDLISLVEATNNFSHENKLGEGGFGSVYKGTLKNGQEIAVKRLSETSKQGWE

EFKNEAKLIAKHTHRNLVRLLGCCAQQEEKLLIYEFMPNKSLDSLIFDKTKNSCLDWKQRFSIIMGIVRGMVYLHHDCSVRIIHR

DLKASNVLLDAQMNPKISDFGMARFFEGDQVEGTTNRVVGTYGYMSPEYAADGTFSIKSDVFSFGVLLLEIITGTKNRSYYND

DEHLNLISKVWNLWREDRALEIVDSSLGGDYSANTVLRCIHVGLLSLQAHAIHRPTMSDIAVMLSYETDLPTPREPGFYFSNFA

SDQHASSSTVGMHSVNEITLSVIEPR

**Figure S5. (A)** Schematic of predicted domain organization of *PdPIRK.* Colored boxes indicate protein motif annotations based on predictions by DeepTMHMM[1] and SIB Swiss Motif Scan My Hits[2] **(B)** Amino acid sequence of *PdPIRK* allele *S4* colored by predicted domain. Domain organization and structure is consistent with the G-type subclass of LecRLKs. These genes contain extracellular bulb lectin, S-locus glycoprotein, and Plasminogen/Apple/Nematode (PAN) domains with a single transmembrane (TM) region and an intracellular kinase domain[3].

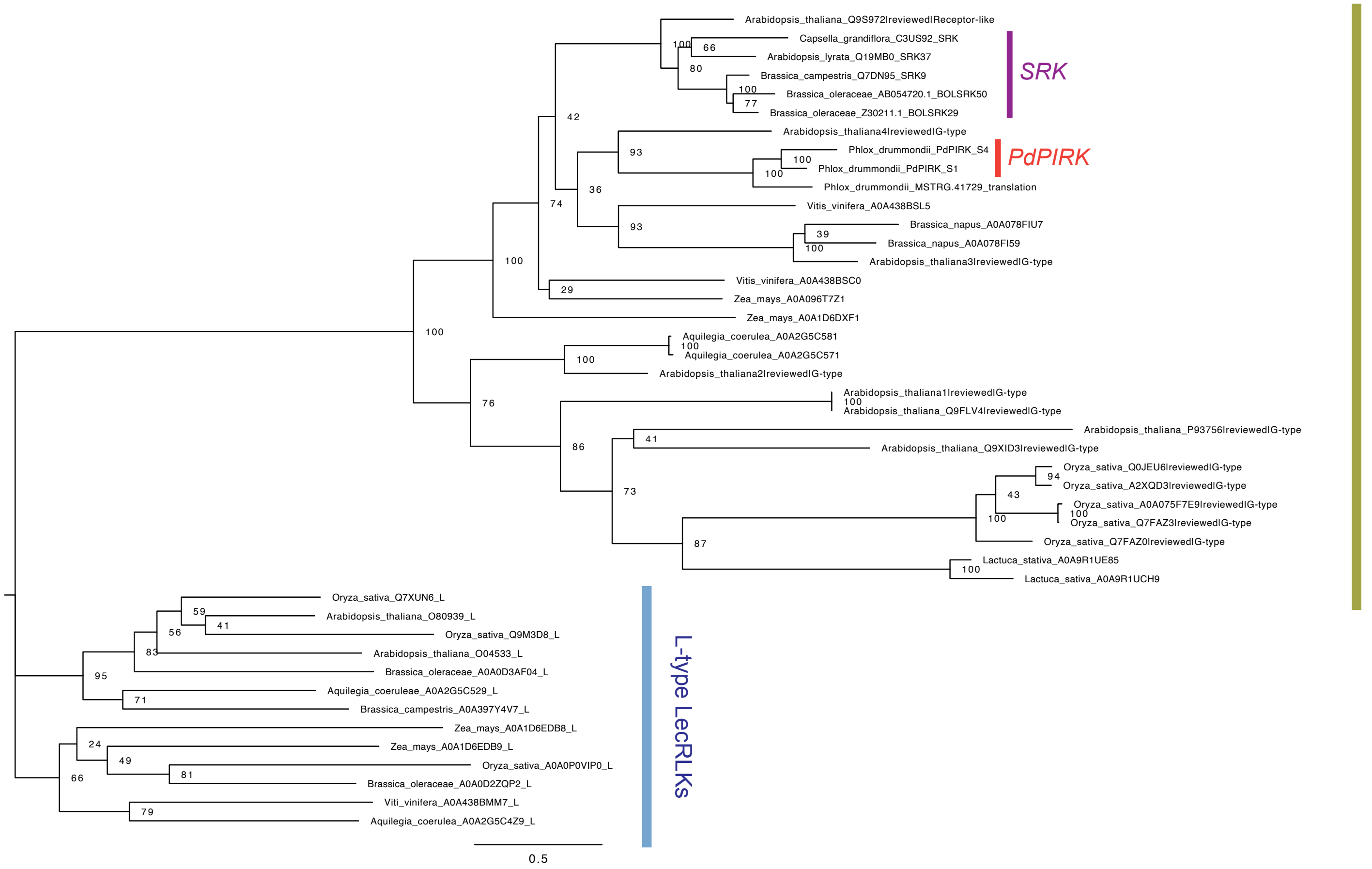


**Figure S6.** Gene tree of G-type and L-type lectin receptor-like kinases. *PdPIRK* is highlighted orange and known sequences of *SRK* are highlighted purple. The tree was generated using RAxML on aligned protein sequences. Bootstrap support is shown for each node. Sequences from *Zea mays, Arabidopsis lyrata, Arabidopsis thaliana, Latuca sativa, Brassica rapa, Brassica napus, Brassica oleraceae, Capsella grandiflora, Aquilegia coerulea,* and *Vitis vinifera* genomes were included.
