## Supplemental Tables for "A lectin receptor-like kinase controls self-pollen recognition in *Phlox* (Polemoniaceae)"

**Table S1.** Genes differentially expressed in stigma/style tissue within the *S*-locus associated region

| Genome Annotation ID | log(Fold Change) | F | p-value | q-value | Contrast |
|---|---|---|---|---|---|
| *Pd51265* | 9.600251665 | 85.9129689 | 5.404E-07 | 8.1166E-06 | stigma/style vs. anther |
| *Pd51265* | 9.600251665 | 85.9129689 | 5.404E-07 | 8.1166E-06 | stigma/style vs. pollen |
| *Pd51265* | 7.693559757 | 69.16110675 | 1.9334E-06 | 3.58631E-05 | stigma/style vs. leaf |
| *Pd51265* | 7.693559757 | 69.16110675 | 1.9334E-06 | 3.58631E-05 | stigma/style vs. stem |
| *Pd51830* | 8.121538088 | 105.4251684 | 1.028E-07 | 2.4768E-06 | stigma/style vs. anther |
| *Pd51830* | 8.121538088 | 105.4251684 | 1.028E-07 | 2.4768E-06 | stigma/style vs. pollen |
| *Pd51830* | 6.174632711 | 57.09017973 | 1.59357E-05 | 0.000152725 | stigma/style vs. leaf |
| *Pd51830* | 6.174632711 | 57.09017973 | 1.59357E-05 | 0.000152725 | stigma/style vs. stem |
| *Pd52359* | 6.082616028 | 28.06576115 | 0.000298739 | 0.000815851 | stigma/style vs. anther |
| *Pd52359* | 6.082616028 | 28.06576115 | 0.000298739 | 0.000815851 | stigma/style vs. pollen |
| *Pd52359* | 5.880994975 | 235.5303958 | 2.96E-08 | 2.5802E-06 | stigma/style vs. leaf |
| *Pd52359* | 5.880994975 | 235.5303958 | 2.96E-08 | 2.5802E-06 | stigma/style vs. stem |
| ***Pd52568*** | **16.28507085** | **277.8747021** | **2.29E-08** | **7.906E-07** | **stigma/style vs. anther** |
| ***Pd52568*** | **16.28507085** | **277.8747021** | **2.29E-08** | **7.906E-07** | **stigma/style vs. pollen** |
| ***Pd52568*** | **13.5126048** | **461.6561342** | **0** | **9.2E-09** | **stigma/style vs. leaf** |
| ***Pd52568*** | **13.5126048** | **461.6561342** | **0** | **9.2E-09** | **stigma/style vs. stem** |
| *Pd52576* | 15.67809738 | 44.97626689 | 0.000116723 | 0.00038594 | stigma/style vs. anther |
| *Pd52576* | 15.67809738 | 44.97626689 | 0.000116723 | 0.00038594 | stigma/style vs. pollen |
| *Pd52576* | 12.80788024 | 41.00776874 | 0.00014801 | 0.000764233 | stigma/style vs. leaf |
| *Pd52576* | 12.80788024 | 41.00776874 | 0.00014801 | 0.000764233 | stigma/style vs. stem |
| *Pd52649* | 8.295679002 | 50.76191108 | 1.80505E-05 | 9.42669E-05 | stigma/style vs. anther |
| *Pd52649* | 8.295679002 | 50.76191108 | 1.80505E-05 | 9.42669E-05 | stigma/style vs. pollen |
| *Pd52649* | 6.846926633 | 62.00249868 | 2.2247E-06 | 3.90377E-05 | stigma/style vs. leaf |
| *Pd52649* | 6.846926633 | 62.00249868 | 2.2247E-06 | 3.90377E-05 | stigma/style vs. stem |
| *Pd52651* | 14.51944186 | 341.6023573 | 3.85E-08 | 1.1744E-06 | stigma/style vs. anther |
| *Pd52651* | 14.51944186 | 341.6023573 | 3.85E-08 | 1.1744E-06 | stigma/style vs. pollen |
| *Pd52651* | 8.659705072 | 225.1673596 | 1.1E-09 | 3.242E-07 | stigma/style vs. leaf |
| *Pd52651* | 8.659705072 | 225.1673596 | 1.1E-09 | 3.242E-07 | stigma/style vs. stem |
| *Pd52670* | 5.23707408 | 35.86351495 | 0.000112291 | 0.000374578 | stigma/style vs. anther |
| *Pd52670* | 5.23707408 | 35.86351495 | 0.000112291 | 0.000374578 | stigma/style vs. pollen |
| *Pd52670* | 7.895285454 | 82.48572411 | 2.3846E-06 | 4.09642E-05 | stigma/style vs. leaf |
| *Pd52670* | 7.895285454 | 82.48572411 | 2.3846E-06 | 4.09642E-05 | stigma/style vs. stem |

**Note.** Gene IDs are given based on the *P. drummondii* genome annotation (NCBI BioProject: PRJNA1219593, unpublished). The annotation ID assigned to *PdPIRK* (*Pd52568*) is bolded and shown in blue.

**Table S2.** Expression and gene tree branch length for the top 10% of genes most highly expressed in stigma/style tissue within the S-locus associated region

| Gene ID | Sample ID (stigma/style) | | | | | | Average Expression | Total Branch Length |
|---|---|---|---|---|---|---|---|---|
| | A5 | C1 | C36 | I6 | I32 | I20 | | |
| *Pd51468* | 3323 | 4019 | 20444 | 14901 | 21531 | 20775 | 14165.5 | 0.021777 |
| *Pd51494* | 6039 | 4347 | 7866 | 17185 | 22547 | 29215 | 14533.16667 | 0.012973 |
| *Pd51500* | 845 | 3132 | 24119 | 3667 | 1650 | 3733 | 6191 | 0.048166 |
| *Pd51508* | 4129 | 2506 | 10706 | 12115 | 7760 | 10818 | 8005.666667 | 0.022797 |
| *Pd51519* | 1447 | 1885 | 5157 | 5033 | 4933 | 5268 | 3953.833333 | 0.017268 |
| *Pd51528* | 3235 | 3800 | 9741 | 10713 | 5126 | 8608 | 6870.5 | 0.144258 |
| *Pd51539* | 5288 | 3098 | 1023 | 16414 | 10346 | 7180 | 7224.833333 | 0.133066 |
| *Pd51552* | 1910 | 2698 | 7005 | 7104 | 8241 | 10529 | 6247.833333 | 0.040551 |
| *Pd51554* | 2116 | 2757 | 7268 | 7124 | 5123 | 8557 | 5490.833333 | 0.034617 |
| *Pd51569* | 1402 | 1642 | 4413 | 5878 | 5179 | 6713 | 4204.5 | 0.022673 |
| *Pd51590* | 2322 | 1899 | 5742 | 5582 | 5936 | 3283 | 4127.333333 | 0.008788 |
| *Pd51610* | 12393 | 10089 | 19447 | 36076 | 18895 | 33688 | 21764.66667 | 2.1e-05 |
| *Pd51624* | 1633 | 3829 | 4391 | 6728 | 6486 | 10760 | 5637.833333 | 0.049696 |
| *Pd51630* | 5218 | 7644 | 10007 | 14688 | 14526 | 10777 | 10476.66667 | 0.114108 |
| *Pd51685* | 4469 | 4219 | 15175 | 13074 | 14584 | 14882 | 11067.16667 | 0.028375 |
| *Pd51703* | 2164 | 4011 | 6379 | 8204 | 6223 | 7220 | 5700.166667 | 0.008026 |
| *Pd51730* | 533 | 1656 | 3336 | 11279 | 4302 | 2593 | 3949.833333 | 0.0023 |
| *Pd51736* | 2486 | 2145 | 5835 | 9344 | 8563 | 5412 | 5630.833333 | 0.014307 |
| *Pd51738* | 3586 | 3421 | 5192 | 6891 | 7564 | 6517 | 5528.5 | 0.007272 |
| *Pd51751* | 1709 | 1875 | 4833 | 5610 | 5095 | 6215 | 4222.833333 | 0.114953 |
| *Pd51788* | 1764 | 2241 | 6580 | 4343 | 5350 | 6725 | 4500.5 | 0.024789 |
| *Pd51794* | 1656 | 2497 | 4490 | 10780 | 9875 | 6506 | 5967.333333 | 0.024227 |
| *Pd51800* | 15999 | 6544 | 32276 | 29511 | 38177 | 40347 | 27142.33333 | 2.1e-05 |
| *Pd51802* | 2722 | 3101 | 8787 | 6700 | 8201 | 9310 | 6470.166667 | 0.002737 |
| *Pd51828* | 1408 | 3608 | 3415 | 6848 | 7293 | 5836 | 4734.666667 | 0.171273 |
| *Pd51866* | 1707 | 2075 | 6272 | 4753 | 7487 | 6867 | 4860.166667 | 0.062473 |
| *Pd51923* | 5003 | 4858 | 12290 | 14990 | 14210 | 14992 | 11057.16667 | 0.00794 |
| *Pd51928* | 42984 | 39956 | 49462 | 95782 | 77817 | 84474 | 65079.16667 | 0.020279 |
| *Pd51941* | 2241 | 2097 | 10783 | 11597 | 11285 | 7961 | 7660.666667 | 0.013468 |
| *Pd51958* | 1979 | 2781 | 7133 | 6087 | 9314 | 6460 | 5625.666667 | 0.014007 |
| *Pd51965* | 2028 | 1830 | 6484 | 5766 | 5655 | 5364 | 4521.166667 | 0.118237 |
| *Pd51970* | 1775 | 1820 | 5061 | 4313 | 5295 | 5142 | 3901 | 0.004929 |
| *Pd51982* | 2119 | 2464 | 7004 | 7432 | 6371 | 7720 | 5518.333333 | 0.008288 |
| *Pd52012* | 3735 | 1748 | 8669 | 9440 | 7987 | 7782 | 6560.166667 | 0.015903 |
| *Pd52036* | 813 | 171 | 1405 | 370 | 339 | 31111 | 5701.5 | 0.025445 |
| *Pd52120* | 1901 | 1329 | 11912 | 4270 | 4847 | 12555 | 6135.666667 | 0.049432 |

| | | | | | | | | |
|---|---|---|---|---|---|---|---|---|
| *Pd52139* | 11285 | 11252 | 41790 | 42211 | 34636 | 66357 | 34588.5 | 0.046286 |
| *Pd52141* | 2759 | 2286 | 4567 | 7749 | 5412 | 7247 | 5003.333333 | 0.012714 |
| *Pd52157* | 18593 | 27948 | 100371 | 72700 | 61046 | 66682 | 57890 | 0.021146 |
| *Pd52182* | 7297 | 9366 | 23990 | 21913 | 23208 | 28679 | 19075.5 | 0.039635 |
| *Pd52228* | 3906 | 2897 | 13659 | 7087 | 7769 | 12871 | 8031.5 | 0.017451 |
| *Pd52230* | 7635 | 6185 | 20767 | 21359 | 16977 | 16453 | 14896 | 0.095145 |
| *Pd52263* | 2573 | 1814 | 5609 | 6050 | 6086 | 5939 | 4678.5 | 0.048981 |
| *Pd52270* | 2452 | 1446 | 4109 | 6192 | 5190 | 5692 | 4180.166667 | 0.01097 |
| *Pd52287* | 1406 | 3789 | 2246 | 5170 | 5344 | 7030 | 4164.166667 | 0.031947 |
| *Pd52317* | 2464 | 1595 | 9577 | 8845 | 10557 | 10887 | 7320.833333 | 0.105934 |
| *Pd52356* | 6890 | 3262 | 10272 | 21185 | 4239 | 8618 | 9077.666667 | 0.136489 |
| *Pd52359* | 40461 | 31448 | 164193 | 79981 | 83805 | 113442 | 85555 | 0.028431 |
| *Pd52464* | 2575 | 2297 | 7251 | 7367 | 7271 | 6854 | 5602.5 | 0.016048 |
| *Pd52466* | 1542 | 3432 | 7011 | 7243 | 5933 | 3549 | 4785 | 0.062661 |
| *Pd52558* | 1972 | 1668 | 5011 | 5607 | 6051 | 5581 | 4315 | 0.066713 |
| ***Pd52568*** | **68403** | **36188** | **178702** | **92859** | **99163** | **75564** | **91813.16667** | **0.296247** |
| *Pd52576* | 2339 | 2165 | 4008 | 34985 | 1332 | 13741 | 9761.666667 | 0.131585 |
| *Pd52613* | 1869 | 1670 | 5806 | 7718 | 4576 | 7380 | 4836.5 | 0.006097 |
| *Pd52641* | 5556 | 4965 | 23421 | 19188 | 29883 | 23598 | 17768.5 | 0.052115 |
| *Pd52646* | 3666 | 8599 | 25160 | 20509 | 19198 | 29022 | 17692.33333 | 0.024163 |
| *Pd52651* | 1189 | 1761 | 5445 | 6070 | 4931 | 4235 | 3938.5 | 0.115151 |
| *Pd52670* | 922 | 2599 | 4537 | 5177 | 13669 | 15364 | 7044.666667 | 0.049441 |
| *Pd52687* | 7820 | 9387 | 23310 | 21990 | 13077 | 32319 | 17983.83333 | 0.166402 |
| *Pd52726* | 1935 | 1030 | 9839 | 3856 | 4455 | 5089 | 4367.333333 | 0.009391 |
| *Pd52753* | 2181 | 3334 | 8777 | 6392 | 8113 | 6781 | 5929.666667 | 0.03042 |
| *Pd52754* | 1379 | 1239 | 5894 | 5456 | 3358 | 6501 | 3971.166667 | 0.007582 |
| *Pd52791* | 1506 | 3428 | 8564 | 4117 | 4997 | 8428 | 5173.333333 | 0.056516 |
| *Pd52793* | 3633 | 1057 | 9783 | 7259 | 3086 | 29637 | 9075.833333 | 0.030442 |
| *Pd52828* | 4112 | 4886 | 12999 | 12335 | 13376 | 10692 | 9733.333333 | 0.056631 |
| *Pd52830* | 1458 | 1529 | 6372 | 6753 | 6468 | 6457 | 4839.5 | 0.052909 |
| *Pd52845* | 1461 | 1599 | 6349 | 5007 | 5750 | 5824 | 4331.666667 | 0.013324 |
| *Pd52879* | 2503 | 917 | 3091 | 4898 | 2744 | 12884 | 4506.166667 | 0.012341 |
| *Pd52933* | 3952 | 1924 | 15670 | 6410 | 8156 | 8625 | 7456.166667 | 0.017508 |
| *Pd52936* | 3433 | 7033 | 10138 | 16923 | 9381 | 7224 | 9022 | 0.050725 |
| *Pd53000* | 3309 | 3292 | 5863 | 7951 | 4694 | 9920 | 5838.166667 | 0.094207 |
| *Pd53001* | 3712 | 4276 | 18560 | 16288 | 11047 | 46296 | 16696.5 | 0.007027 |
| *Pd53012* | 4787 | 1725 | 3684 | 8856 | 4642 | 4469 | 4693.833333 | 0.011807 |

**Note.** Expression is given in TMM, total gene tree branch length is given in substitutions per site. The *P. drummondii* genome annotation ID assigned to *PdPIRK* (*Pd52568*) is bolded and shown in blue.

**Table S3.** Primers used for PCR-genotyping and RT-qPCR of *PdPIRK*

| Allele | Forward | Reverse | Annealing (°C) |
|---|---|---|---|
| *S1* | TTTACCAAGGGTCGAATCGGATATG | GTAATCGTCACAATACCATTTGGGC | 66.5 |
| *S2* | GTTTCAGAAAACGGAATTGTTTGGG | ATATTTGGAAGCCAAGTGTCTGTTG | 65.5 |
| *S3* | ATATCATCAATGCCCAGCTTCAATG | GACCAGAGCACAATATTTGTTTGGA | 65.5 |
| *S4* | GGACTTGACCTGAAAATCCCAATTT | GGGTCATCAATTGATTTCCAGGATC | 66.5 |

**Table S4.** Pre-treatment phenotypes for plants used in VIGS experiments

| Individual | *PdPIRK* Genotype | VIGS TRV2 Treatment | Proportion Viable Pollen | Outcross Fruit (out of 8) |
|---|---|---|---|---|
| vA_40 | *S4S4* | *PdPIRK(S4)-DFR* | 0.959183673 | 8 |
| vA_44 | *S4S4* | *PdPIRK(S4)-DFR* | 0.950980392 | 8 |
| vA_63 | *S4S4* | *PdPIRK(S4)-DFR* | 0.994736842 | 8 |
| vA_72 | *S4S4* | *PdPIRK(S4)-DFR* | 0.989361702 | 8 |
| vA_76 | *S4S4* | *PdPIRK(S4)-DFR* | 0.948979592 | 8 |
| vA_31 | *S2S4* | *PdPIRK(S4)-DFR* | 0.964285714 | 8 |
| vA_33 | *S2S4* | *PdPIRK(S4)-DFR* | 0.984375 | 8 |
| vA_36 | *S2S4* | *PdPIRK(S4)-DFR* | 0.987951807 | 6 |
| vA_56 | *S2S4* | *PdPIRK(S4)-DFR* | 0.95890411 | 6 |
| vA_50 | *S2S4* | *PdPIRK(S4)-DFR* | 0.970238095 | 8 |
| vA_105 | *S1S1* | *PdPIRK(S1)-DFR* | 0.9090909091 | 8 |
| vA_116 | *S1S1* | *PdPIRK(S1)-DFR* | 0.9574468085 | 8 |
| vA_119 | *S1S1* | *PdPIRK(S1)-DFR* | 0.9453551913 | 8 |
| vA_123 | *S1S1* | *PdPIRK(S1)-DFR* | 0.9069767442 | 5 |
| vA_101 | *S1S4* | *PdPIRK(S1)-DFR* | 0.9277108434 | 8 |
| vA_103 | *S1S4* | *PdPIRK(S1)-DFR* | 0.95 | 7 |
| vA_112 | *S1S4* | *PdPIRK(S1)-DFR* | 0.9827586207 | 6 |
| vA_107 | *S4S4* | untreated pollen donor | 0.975308642 | NA |
| vA_34 | *S2S2* | untreated pollen donor | 0.96 | NA |
| vA_85 | *S2S2* | untreated pollen donor | 0.9375 | NA |
| vA_1 | *S4S4* | *DFR* | 0.919708029 | 8 |
| vA_2 | *S2S4* | *DFR* | 0.996666667 | 8 |
| vA_3 | *S2S2* | *DFR* | 1 | 8 |
| vA_8 | *S2S4* | *DFR* | 0.989583333 | 8 |
| vA_10 | *S2S2* | *DFR* | 0.968208092 | 8 |
| vA_11 | *S1S4* | *DFR* | 0.987951807 | 8 |

**Table S5.** Sequences inserted into TRV2 silencing vector

| Gene Target | Insert Sequence |
|---|---|
| *DFR* | GAACGTTGAAGAGCATCAAAAACCCGTTTACAACGAGAACAATTGGAGCGACTTGGATTTTATCT<br>ACAGCAAGAAAATGACTGCCTGGATGTATTTTGTGTCAAAGACATTGGCAGAGAAAGCAGCATG<br>GGAAGCTGCTAAAGAAAACAAGATTGATTTCATTAGCATCATACCACCATTAGTGGTTGGTCCGT<br>TCATTACTCCAACATTCCCACCTAGCCTAATCACAGCCCTCTCTCCCATCACCGGAAACGAAGC<br>GCACTATTCAATCATAAAGCAAGGGCAATTTGTGCACCTGGA |
| *PdPIRK(S4)* | ACCTCGATGGCGGGGGGTCTGGAAGGCCCCCGAGGGCCTTTGTGACCACTATGGCATGTGTG<br>GTCCGAATCGAAACTGTGATGCTGGCAGTACAGCGCCTTTGATCAATTGCGAATGTTTGCCCGG<br>GTCAAAAATCAAGTCTCAACGTGAATTTGATGCCGGAAACATGTCAGGAGGTTGTGTACGGAAC<br>CGTGACCGTAATGATTGTTGGAACGGAGAAGGGTT |
| *PdPIRK(S1)* | GACTCTGCCCAAATGGTATTGTGACGATTACAATCGCTGTGGTGAGAACAAATATTGTGCTGCT<br>CTCACTGAAACAGTCACGTCTGAGTGCTCATGCTTTCCTGGATTTGAACCGAAAAACATAAAGGT<br>AGTTGGAGGAGGGTGCGTGAGGAAGCACGGCCGTAACGTTTGTCGAAACGGCGAAGGGTTTG<br>CCAAATTTTCAAATATGAAGATGCCTGACACTAC |

**Table S6.** NCBI accession numbers for sequences used to construct a gene tree of G-type and L-type lectin receptor-like kinases (Figure S6)

| NCBI Accession | Sequence ID | Species |
|---|---|---|
| PIA26311.1 | Aquilegia_coerulea_A0A2G5C4Z9_L | *Aquilegia coerulea* |
| PIA26432.1 | Aquilegia_coerulea_A0A2G5C571 | *Aquilegia coerulea* |
| PIA26426.1 | Aquilegia_coerulea_A0A2G5C581 | *Aquilegia coerulea* |
| PIA26376.1 | Aquilegia_coeruleae_A0A2G5C529_L | *Aquilegia coerulea* |
| ABF71379.1 | Arabidopsis_lyrata_Q19MB0_SRK37[1] | *Arabidopsis lyrata* |
| NP_177170.1 | Arabidopsis_thaliana_O04533_L | *Arabidopsis thaliana* |
| NP_181307.1 | Arabidopsis_thaliana_O80939_L | *Arabidopsis thaliana* |
| NP_001318404.1 | Arabidopsis_thaliana_P93756\|reviewed\|G-type | *Arabidopsis thaliana* |
| NP_001330805.1 | Arabidopsis_thaliana_Q9FLV4\|reviewed\|G-type | *Arabidopsis thaliana* |
| NP_176756.1 | Arabidopsis_thaliana_Q9S972\|reviewed\|Receptor-like | *Arabidopsis thaliana* |
| NP_174690.1 | Arabidopsis_thaliana_Q9XID3\|reviewed\|G-type | *Arabidopsis thaliana* |
| NP_849636.1 | Arabidopsis_thaliana1\|reviewed\|G-type | *Arabidopsis thaliana* |
| NP_179503.1 | Arabidopsis_thaliana2\|reviewed\|G-type | *Arabidopsis thaliana* |
| NP_563887.1 | Arabidopsis_thaliana3\|reviewed\|G-type | *Arabidopsis thaliana* |
| NP_172601.3 | Arabidopsis_thaliana4\|reviewed\|G-type | *Arabidopsis thaliana* |
| XP_033135722.1 | Brassica_campestris_A0A397Y4V7_L | *Brassica rapa* |
| BAA21132.1 | Brassica_campestris_Q7DN95_SRK9[2] | *Brassica rapa* |
| XP_013717485.2 | Brassica_napus_A0A078FI59 | *Brassica napus* |
| CDY12902.1 | Brassica_napus_A0A078FIU7 | *Brassica napus* |
| KAL0669094.1 | Brassica_oleraceae_A0A0D2ZQP2_L | *Brassica oleraceae* |
| XP_013616924.1 | Brassica_oleraceae_A0A0D3AF04_L | *Brassica oleraceae* |
| AB054720.1 | Brassica_oleraceae_AB054720.1_BOLSRK50[3] | *Brassica oleraceae* |
| Z30211.1 | Brassica_oleraceae_Z30211.1_BOLSRK29[3] | *Brassica oleraceae* |
| ACO83289.1 | Capsella_grandiflora_C3US92_SRK[3] | *Capsella grandiflora* |
| KAJ0184612.1 | Lactuca_sativa_A0A9R1UCH9 | *Lactuca sativa* |
| KAJ0185460.1 | Lactuca_stativa_A0A9R1UE85 | *Lactuca sativa* |

| | | |
|---|---|---|
| NP_001403531.1 | Oryza_sativa_A0A0P0VIP0_L | *Oryza sativa* |
| NP_001406034.1 | Oryza_sativa_A0A075F7E9\|reviewed\|G-type | *Oryza sativa* |
| Swiss-Prot: A2XQD3.1 | Oryza_sativa_A2XQD3\|reviewed\|G-type | *Oryza sativa* |
| XP_025880371.1 | Oryza_sativa_Q0JEU6\|reviewed\|G-type | *Oryza sativa* |
| AIE56246.1 | Oryza_sativa_Q7FAZ0\|reviewed\|G-type | *Oryza sativa* |
| AIE56229.1 | Oryza_sativa_Q7FAZ3\|reviewed\|G-type | *Oryza sativa* |
| XP_015635472.1 | Oryza_sativa_Q7XUN6_L | *Oryza sativa* |
| NP_190127.1 | Oryza_sativa_Q9M3D8_L | *Oryza sativa* |
| RVW12226.1 | Viti_vinifera_A0A438BMM7_L | *Vitis vinifera* |
| RVW13862.1 | Vitis_vinifera_A0A438BSC0 | *Vitis vinifera* |
| RVW13860.1 | Vitis_vinifera_A0A438BSL5 | *Vitis vinifera* |
| NP_001145772.2 | Zea_mays_A0A1D6DXF1 | *Zea mays* |
| ONM18285.1 | Zea_mays_A0A1D6EDB8_L | *Zea mays* |
| ONM18286.1 | Zea_mays_A0A1D6EDB9_L | *Zea mays* |
| XP_008674953.1 | Zea_mays_A0A096T7Z1 | *Zea mays* |

**Note.** Lectin domain containing receptor-like kinase sequences were identified using InterPro (G-type LecRLKs: IPR024171; L-type LecRLKs: IPR050528). Sequence ID is provided as it appears in Figure S6. References are given for previously characterized *SRK* sequences.
